# Revision of Archaeosporomycetes with two old and two new fungal orders: Archaeosporales, Geosiphonales, Polonosporales, and Ambisporales

**DOI:** 10.64898/2026.03.05.709871

**Authors:** Fritz Oehl, Janusz Błaszkowski, Ewald Sieverding, Piotr Niezgoda, Thays Gabrielle Lins de Oliveira, Daniele Magna Azevedo de Assis, Viviane Monique Santos, Bruno Tomio Goto, Mike Anderson Corazon-Guivin, Gladstone Alves da Silva

## Abstract

Currently, the fungal class Archaeosporomycetes consists of one order, Archaeosporales with four families: Archaeosporaceae, Ambisporaceae, Geosiphonaceae, and Polonosporaceae. In the present study, the objective was to re-analyze the phylogeny and morphology of the Archaeosporomycetes from order to genus level. The different ecological strategies and, consequently, distinct evolutionary patterns of these taxa, as well as their morphological characters and other data updated here, suggest the need to divide Archaeosporales into four orders: (i) the type order Archaeosporales, (ii) Ambisporales ord. nov., both with four genera, (iii) Geosiphonales and (iv) Polonosporales ord. nov., both with single families and genera. Remarkably, the order Geosiphonales was described in the past, but was not considered in the Archaeosporomycetes until now. Phylogenetically, the four main clades (orders here proposed) of Archaeosporomycetes are well supported, with bootstrap values higher than 95% in all analyses, except Ambisporales/Ambisporaceae for RAxML-NG FBP analysis in the SSU tree (75%). Ecologically, this class includes three orders of arbuscular mycorrhizal fungi (AMF) forming symbiotic associations with plants, while Geosiphonales form an endocytobiosis with the cyanobacterium *Nostoc*. Morphologically, there are at least two AMF orders with spore bimorphism, which has not (yet) been described for Polonosporales. The only known species of Polonosporales, *Polonospora polonica*, forms spores directly on the neck of sporiferous saccules and the spores can morphologically be differentiated from all other taxa in Archaeosporomycetes by the formation of three permanent, rather thick spore walls, of which two form *de novo* during spore formation. The outer spore wall of Archaeosporales and Ambisporales are semi-permanent, evanescent or even short-lived, or show multiple fissures during aging, when it is more resistant. Ambisporales can easily be differentiated from Archaeosporales for instance by larger spores of the acaulosporoid morph and thicker spore walls. Our phylogenetic analyses suggested that Archaeosporales can be divided into two families: Antiquisporaceae that was described to form intraradical hyphae, vesicles and spores, staining darkly in Trypan blue, and Archaeosporaceae whose hyphae generally do not or only faintly stain in this reagent, and vesicles and intraradical spores have been rarely, if ever reported.

## Introduction

The fungal class Archaeosporomycetes was separated from Glomeromycetes together with the class Paraglomeromycetes following concomitant phylogenetic and morphological analyses (Oehl et al. 2011a; Hyde et al. 2024; Wijayawardene et al. 2025). The large class Glomeromycetes so far counts seven fungal orders, i.e. Glomerales, Acaulosporales Diversisporales, Entrophosporales, Gigasporales, Pacisporales, and Sacculosporales (Schüßler et al. 2001; Walker and Schüßler 2004; Oehl et al. 2011a; Błaszkowski et al. 2022; Oehl et al. 2026), while the other two classes have remained each with one order only so far. Nevertheless, several genera were attributed to Archaeosporomycetes in the last fifteen years, i.e. beside the type genus *Archaeospora*, also *Intraspora*, *Palaeospora*, *Andinospora*, and *Antiquispora* in the family Archaeosporaceae, while *Ambispora*, the type genus of Ambisporaceae, was grouped with *Appendiculaspora*, *Ephemerapareta*, and *Pelotaspora* (Spain et al. 2006; Walker et al. 2007; Esmaeilzadeh-Salestani et al. 2025; Silva et al. 2026). Geosiphonaceae and Polonosporaceae hitherto form monogeneric families with *Polonospora* and *Geosiphon* as type genera, respectively (Schüßler and Walker 2010; Błaszkowski et al. 2021).

Major morphological differences have well been known between the current orders of Glomeromycetes, which are for instance the formation of typical vesicular-arbuscular mycorrhiza (VAM) in the roots, or formation of arbuscular mycorrhiza (AM) without intraradical vesicles, or the specific type of spore formation (Morton and Benny 1990; Oehl et al. 2011b, c; Silva et al. 2024). These differences led to a separation on the order level, however, clearly driven by the phylogenetic analyses (Oehl et al. 2026).

Within Archaeosporomycetes, such morphological differences are even more evident. In Archaeosporaceae, the spores are usually < 100 µm, hyaline to subhyaline to rarely light creamy, rather thin-walled, and the intraradical hyphae have also a tiny small diameter, vesicles are rarely, if ever observed, and, besides spores, the intraradical structures generally do not stain or only faintly in Trypan blue or ink (Oehl et al. 2019), with the notable exception of *Antiquispora* as described by Esmaeilzadeh-Salestani et al. (2025). In Ambisporaceae, spores are generally > 100 µm, and often > 200 µm, they are usually pigmented quite thick-walled, and the intraradical mycorrhizal and especially the extraradical mycelial hyphae have also rather large diameters and rather thick walls, which generally stain pale blue in Trypan blue or ink (Spain et al. 2006). However, the greatest differences are known for Geosiphonaceae, which does not form symbiosis with plants, but an endocytobiosis with the cyanobacterium *Nostoc* (Gehrig et al. 1996). Thus, it has a completely different life cycle and ecological strategy when compared to all other Glomeromycota taxa, which form typical VAM or AM symbiosis with higher plants, mosses or ferns (Smith and Read 2008; Wijayawardene et al. 2025). Cavalier-Smith (1998) described an order and a specific class (Geomycetes) to *Geosiphon*. This fungus is so distinct that, initially, this author placed it in Ascomycota. Benny et al. (2001) decided to transfer Geosiphonales to former Zygomycetes, based on the morphology of their reproductive structures. At that time, the phylum Glomeromycota had not yet been described, but phylogenetic evidence already indicated that *Geosiphon* was a basal group in Glomerales and their spores resembled those from *Glomus* (Gehrig et al. 1996). However, Benny et al. (2001) considered Geosiphonales as a sister group to Glomerales. In the same year, phylogenetic analyses by Schüßler et al. (2001) demonstrated that Glomerales was in fact a phylum (Glomeromycota) and described the new order Archaeosporales, to which the family Geosiphonaceae was attributed. These authors indicated that the symbiosis formed by *Geosiphon* could demonstrate an initial evolutionary stage in AM-like associations (see also Schüßler 2002). Otherwise, Nelsen et al. (2025) report that *Geosiphon*-like state is a derived condition, being, *Geosiphon*, a taxon inside Glomeromycota, not an ancestor of the entire clade.

The objective of the present study was to thoroughly analyze the current molecular and morphological data sets of Archaeosporomycetes for higher-level differentiation and to summarize the current stage of knowledge. Our hypothesis was that the datasets we had collected clearly suggest that splitting the current order Archaeosporales into two, or even several, orders would be justified. Therefore, in the taxonomic section, this division would be made according to the results of phylogenetic analyses. Finally, we assumed that potential new orders could be divided into several, or even a dozen families, whose descriptions would then be presented in the taxonomic section.

## Materials and Methods

### Phylogenetic analyses

To reconstruct the phylogeny, two alignments (datasets), based on partial SSU, ITS region and partial LSU nrDNA (dataset 1 - SSU+ITS+LSU) and complete SSU nrDNA (dataset 2 - SSU) were generated with AM fungal sequences (Supplementary Material, Spreadsheet S1). *Paraglomus brasilianum* (Spain & J. Miranda) J.B. Morton & D. Redecker was included as outgroup for both datasets. The sequences available for *Geosiphon pyriformis* do not have the ITS1 and ITS2 regions, thus the sequences with the partial SSU and partial LSU nrDNA were concatenated with the only 5.8s sequence and used for the dataset 1. All *Geosiphon pyriformis* sequences used are from the same isolate. The sequences from *Archaeospora europaea* used here have just the partial SSU and ITS nrDNA fragment. The datasets were aligned in Mafft v.7 (Katoh et al. 2019) using the default parameters. Prior to phylogenetic analyses, the model of nucleotide substitution was estimated with ModelTest-NG (Darriba et al. 2020). Maximum Likelihood - ML (1000 bootstrap) analyses were performed using RAxML-NG with Felsenstein Bootstrap Proportion (FBP) and Transfer Bootstrap Expectation (TBE), respectively (Stamatakis 2014; Kozlov et al. 2019; Edler et al. 2020).

### Morphological analyses

Specimens of almost all AMF species currently attributed to Ambisporaceae were available for us (Silva et al. 2026). We morphologically analyzed additionally almost all species of Archaeosporaceae and Polonosporaceae described before 2016, in particular those, which were originally described from our laboratories, i.e. *Intraspora schenckii* (Sieverding and Toro 1987), *Archaeospora undulata* (Sieverding 1988), *Ar. europaea* (Oehl et al. 2019), *Palaeospora spainiae* (Oehl et al. 2015), *Polonospora polonica* (Błaszkowski 1988, re-described by Błaszkowski et al. 2021), and *Antiquispora disseminans* (Esmaeilzadeh-Salestani et al. 2025), but also a specimen of *Ar. trappei* isolated from the type location in Oregon (USA) by Joyce L. Spain (Spain 2003). Unfortunately, we did not have access to *Andinospora ecuadoriana* (Archaeosporales) and *Geosiphon pyriformis* (Geosiphonales). Older specimens (mounted on microscopic slides prior to 1990) were mostly mounted in lactophenol, while others were fixed with polyvinyl alcohol–lactic acid–glycerol (PVLG) or in a mixture of PVLG + Melzer’s reagent, which after 1990 are the principal fixing media (Brundrett et al. 1994). Newly mounted spores and sporocarps from collections or from cultures were fixed using the latter two fixing media or occasionally also a mixture of 1:1 lactic acid to water, Melzer’s reagent and water (Spain 2003). When available, spores freshly isolated from soil or bait cultures were also mounted and analyzed. All spore observations and all information on spore characteristics are based on spores extracted from soil, trap cultures, or single or multiple spore-derived pure cultures. No information is provided from *in vitro* cultured materials. Spore wall terminology follows the nomenclature of Walker (1983) and Błaszkowski (2012). Analyses of the spore walls, germination structures, and all mycorrhizal structures were performed using compound microscopes at 100–1000× magnification. For this paper, all original species descriptions and published species emendations were also considered and thoroughly studied, especially for those species, for which no specimen was available to us.

## Results and discussion

### Molecular phylogeny

The dataset 1 was generated with 78 sequences and 1763 sites; of which 887 were constant, 55 variable (but not parsimony informative), and 821 parsimony informative. For the dataset 2, 57 sequences with 1671 sites were used, of which 1311 were constant, 116 variable (but not parsimony informative), and 244 were parsimony informative. The phylogeny generated by both datasets, using RAxML-NG (FBP) and (TBE), showed similar topology.

Currently, Archaeosporomycetes is represented by the single order Archaeosporales, with four families (Ambisporaceae, Archaeosporaceae, Geosiphonaceae, and Polonosporaceae). In the tree generated by Tedersoo et al. (2024), Geosiphonaceae is in the base of Archaeosporomycetes, but with low support to maintain this family into the class. In other studies, Geosiphonaceae was included in Archaeosporomycetes with strong phylogenetic support (Schüßler et al. 2001; Oehl et al. 2011a, b; Krüger et al. 2012; Wetzel et al. 2014; Błaszkowski et al. 2021; Esmaeilzadeh-Salestani et al. 2025). In our phylogenetic analyses (Figs. 1 and 2), Geosiphonales/Geosiphonaceae is included in Archaeosporomycetes together with other three major clades well supported, all of them with bootstrap values higher than 95% in all analyses, except Ambisporales/Ambisporaceae in the SSU tree (75%) from RAxML-NG FBP analysis. The four major clades represent the four orders proposed in the present study.

**Fig. 1.**
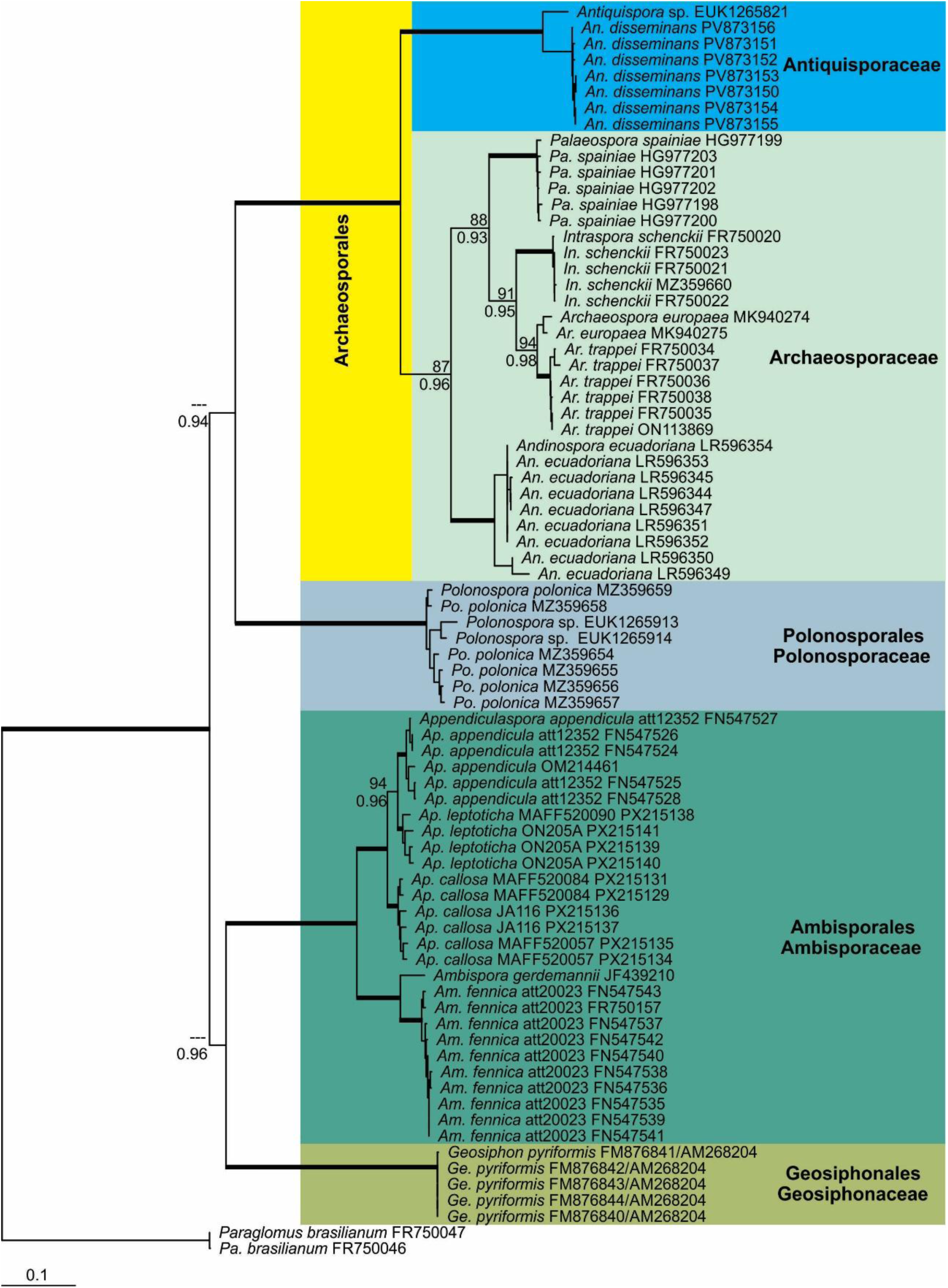
Phylogenetic tree obtained by analysis of partial SSU, ITS region, and partial LSU nrDNA sequences of Archaeosporomycetes. Sequences are labelled with their database accession numbers. Support values (from top) are from ML (maximum likelihood analysis) using RAxML-NG with FBP (Felsenstein Bootstrap Proportion) and TBE (Transfer Bootstrap Expectation), respectively. Only support values of at least 70% are shown. Thick branches represent clades with more than 95% of support in all analyses. The tree was rooted by sequences from *Paraglomus brasilianum*. The nucleotide substitution model used was GTR+I+G for both analyses.

**Fig. 2.**
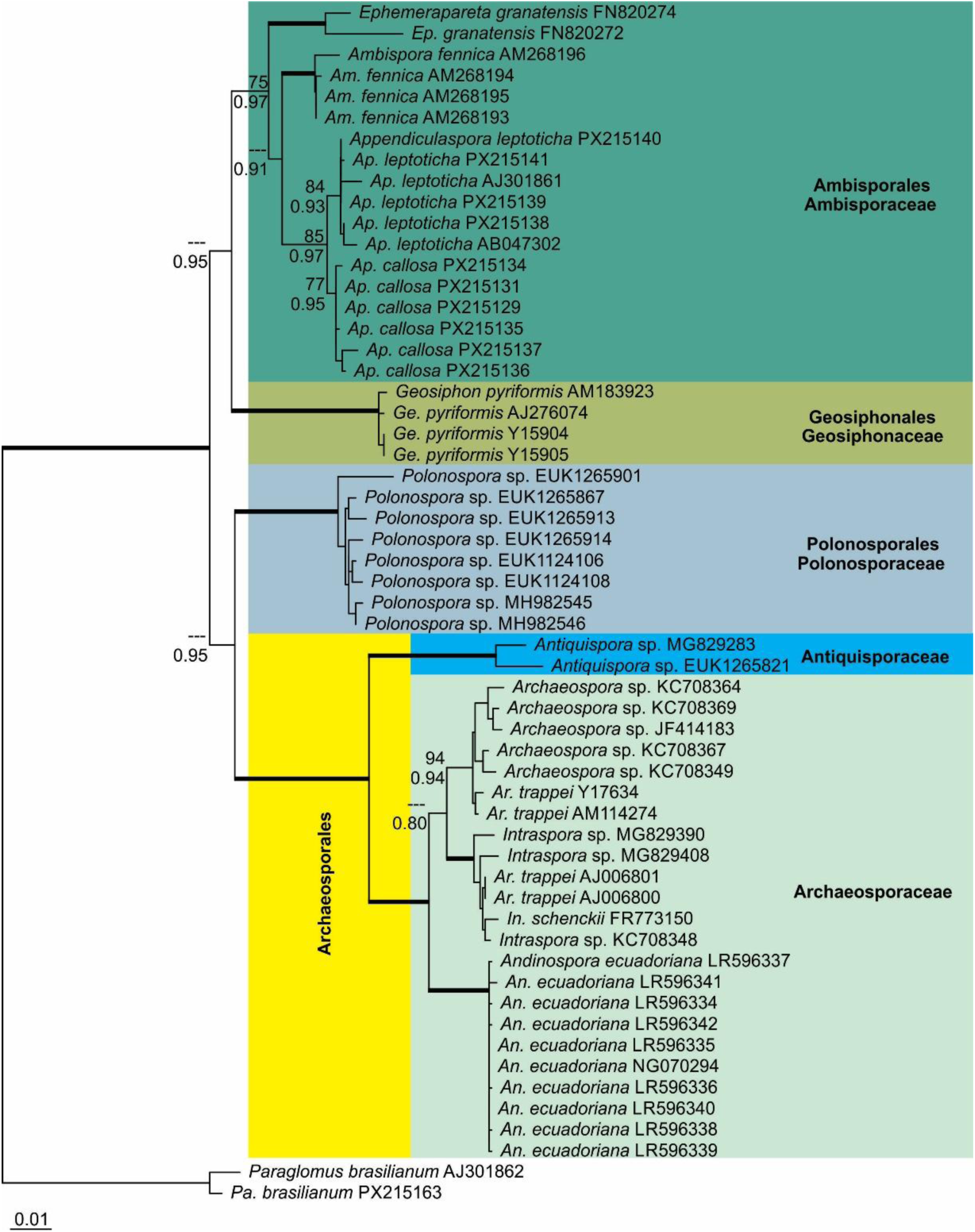
Phylogenetic tree obtained by analysis of complete SSU nrDNA sequences of Archaeosporomycetes. Sequences are labelled with their database accession numbers. Support values (from top) are from ML (maximum likelihood analysis) using RAxML-NG with FBP (Felsenstein Bootstrap Proportion) and TBE (Transfer Bootstrap Expectation), respectively. Only support values of at least 70% are shown. Thick branches represent clades with more than 95% of support in all analyses. The tree was rooted by sequences from *Paraglomus brasilianum*. The nucleotide substitution model used was TIM3+I+G for both analyses.

The genus *Antiquispora* is separated within Archaeosporaceae with bootstrap values > 95% in all analyses (Figs. 1 and 2), suggesting a separation on the family level (hereafter called Antiquisporaceae). All genera in both trees presented support values above 90%, except *Appendiculaspora* in the SSU tree (85%; Fig. 2) from RAxML-NG FBP analysis. The genera not represented in the SSU+ITS+LSU tree were *Ephemerapareta*, and *Pelotaspora* (Fig. 1), because *Ep. granatensis* has only sequences from the ITS region and complete SSU nrDNA, and *Pelotaspora* was described based only on morphological analysis. No environmental sequences from SSU+ITS+LSU were found for *Ephemerapareta*. *Palaeospora*, and *Pelotaspora* also were not represented in the SSU tree (Fig. 2), because *Pelotaspora* was described based just on morphology, and no sequences, not even environmental sequences, are available from complete SSU nrDNA for *Palaeospora.* The genera *Polonospora* and *Antiquispora* were represented just by environmental sequences in the SSU tree (Fig. 2).

The genus *Intraspora*, with the only species *In. schenkii*, was synonymized with *Archaeospora* by Schüßler and Walker (2010). Afterwards, *Archaeospora schenkii* was synonymized with *Archaeospora trappei* by Bills (2015). In our phylogenetic analyses, *Intraspora* is, however, confirmed as a separate clade and thus, independent genus in Archaeosporaceae (Figs. 1 and 2). This genus showed support higher than 95% by all analyses in both trees. Two isolates attributed to *Ar. trappei* are placed in the *Intraspora* clade in the SSU tree, nonetheless, we strongly assume that these isolates are, in fact, from *Intraspora* species. Recently, Esmaeilzadeh-Salestani et al. (2025) found several *Archaeospora* isolates grouping within the clade of a new genus, *Antiquispora*. According to these authors, these *Archaeospora* isolates belong to *Antiquispora*. Futhermore, *Intrapora schenkii* presented about 90% of Maximum Identity (MI) with the nearest species (*Archaeospora trappei*), considering the entire “barcode” fragment (partial SSU + ITS + partial LSU). According to Silva et al. (2023), differences about 10% in MI are indicative for the separation of genera, for instance, in Glomerales.

### Morphological analyses

In this study, no new morphological findings are reported, but the general knowledge collected from the already described families, genera, and species belonging nowadays to the Archaeosporomycetes is summarized in Tab. 1. Certainly, Geosiphonaceae is the most outstanding with its symbiosis between the glomeromycotean fungus *Geosiphon pyriformis* and the cyanobacteria *Nostoc* (Cavalier-Smith 1998; Schüßler 2002; Wijayawardene et al. 2025). Therefore, it is here a logical step to resurrect the order Geosiphonales and transfer it formally from the former Zygomycetes to its family Geosiphonaceae and its type genus *Geosiphon* with the type species *Ge. pyriformis* within Archaeosporomycetes, not only due to its phylogenetic position, but also due to its unique ecological patterns. The *Geosiphon* spores themselves do not have any unique characters, which would deserve a separation on the order level (Schüßler et al. 1994; Schüßler 2002; Wijayawardene et al. 2025), or clear characters which would help to clearly separate them from glomoid spores of Archaeosporaceae or Paraglomeromycetes, except those of the large majority of members of the Glomeromycetes (e.g. Oehl et al. 2011b; Silva et al. 2024; Oehl et al. 2026).

**Tab. 1:**
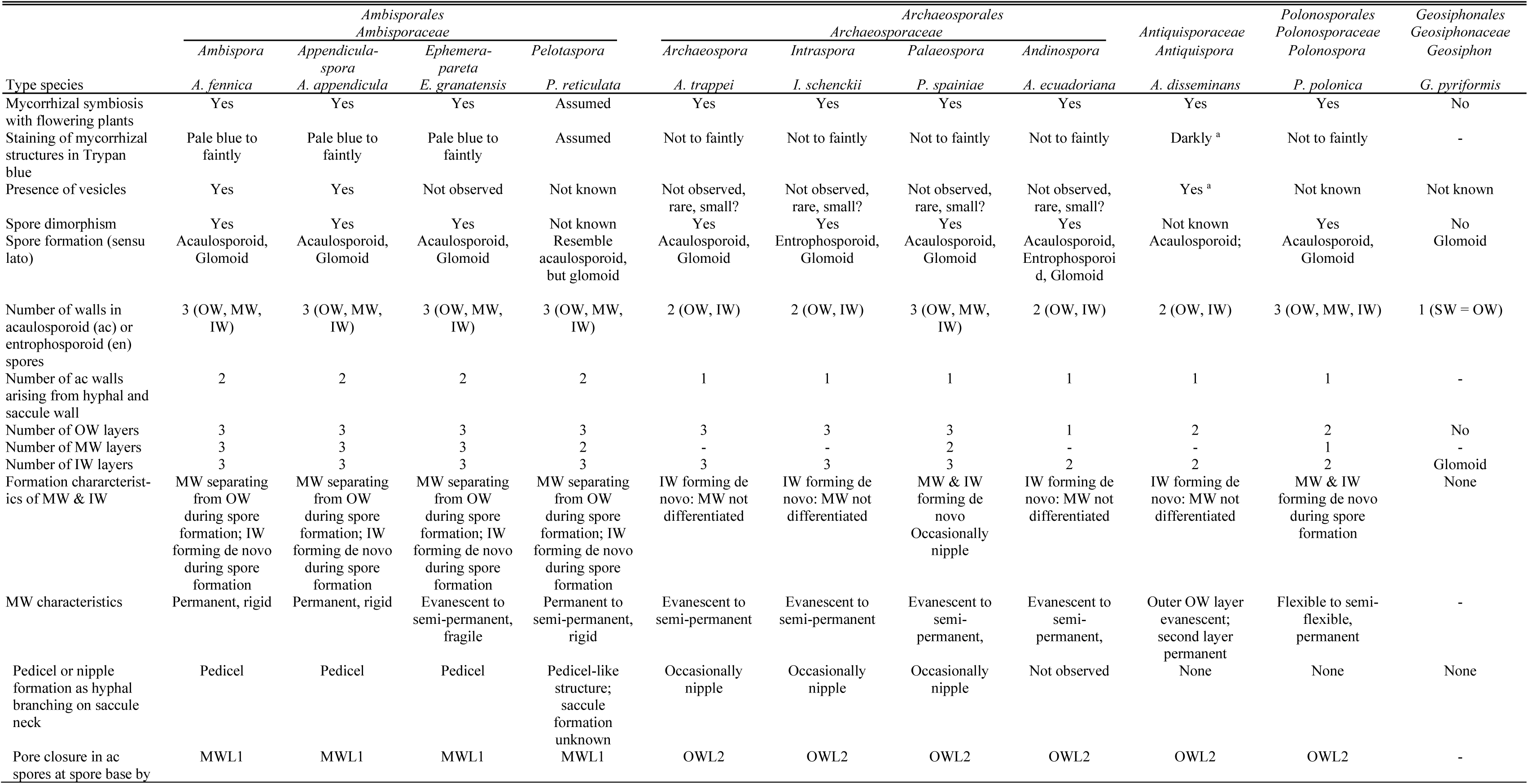
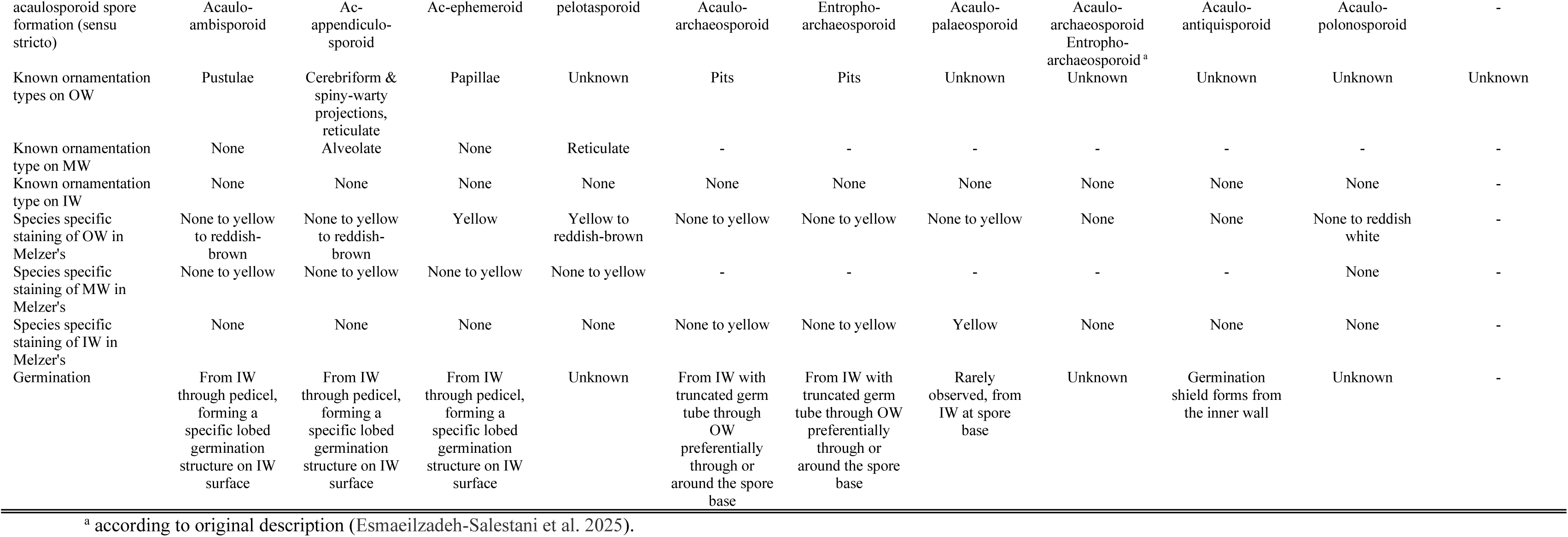
Major morphological characteristics separating genera of Archaeosporomycetes.

Also, a morphological separation between the other families in Archaeosporomycetes, Archaeosporaceae, Ambisporaceae, and Polonosporaceae, can be justified on the order level, following the arguments recently used for the new classification of Glomeromycetes (Oehl et al. 2026). Nevertheless, it should be stressed here that since 2001 all the separations on the order level were always driven clearly by molecular phylogeny (Schüßler et al. 2001, Oehl et al. 2011a, b, c; Błaszkowski et al. 2022; Silva et al. 2024; Silva et al. 2025; Oehl et al. 2026).

In summary, morphologically, there are at least two AMF orders with spore bi-morphism, which has not (yet) been observed for Polonosporales. Ambisporaceae form their acaulo-ambisporoid spores generally on pedicels arising from the neck of sporiferous saccules, with three clearly visible, thick walls (Table 1). Ambisporales can easily be differentiated from Archaeosporales for instance by larger spores of the acaulosporoid morph and thicker spore walls. Polonosporales forms spores directly on the neck of sporiferous saccules. It can morphologically be differentiated from all other taxa in Archaeosporomycetes by the formation of three permanent, rather thick spore walls, of which two form *de novo* during spore formation. In contrast, the outer spore wall of Archaeosporales and Ambisporales is semi-permanent, evanescent or even very short-lived, degrading within weeks, or showing multiple fissures during aging, when they are more resistant. The order Archaeosporales can be divided into two families, as suggested by the phylogenetic analyses presented in this study: Antiquisporaceae that was described to form intraradical hyphae, spores, and vesicles (Esmaeilzadeh-Salestani et al. 2025), which stain darkly in Trypan blue, and Archaeosporaceae whose hyphae generally do not or only faintly stain, and vesicles have been rarely if ever reported.

### Taxonomy

**Archaeosporomycetes** Sieverd., G.A. Silva, B.T. Goto & Oehl, Mycotaxon 116: 374. 2011a; emended here by Oehl, Sieverd, G.A. Silva & Błaszk.

MycoBank no.: MB 519686

Emended description: see below description for the new subclass and Oehl et al. (2011a)

Type subclass: Archaeosporomycetidae Oehl, Sieverd, G.A. Silva, B.T. Goto & Błaszk. subcl, nov.

**Archaeosporomycetidae** Oehl, Sieverd, G.A. Silva, B.T. Goto & Błaszk. subcl. nov.

MycoBank no.: MB 862463

Description: Forms symbiosis between plant roots and fungi, or between cyanobacteria and fungi; if between plants and fungi, fungal spores with two morphs, rarely only one morph or three morphs (ambi-acaulosporoid or archaeo-acaulosporoid, glomoid (sensu lato), archaeo-entrophosporoid) known, and forming vesicular arbuscular or arbuscular mycorrhiza which regularly do not stain or only faintly to pale blue with Trypan blue, and spores, which may stain blue to dark blue.

Etymology: archaios (Greek, old), spora (Greek spore, seed), referring to the ancestral position of this taxon within Glomeromycota.

Type order: Archaeosporales C. Walker & A. Schüssler

Other orders:

Ambisporales Oehl, Sieverd., G.A. Silva & Błaszk.

Geosiphonales Caval.-Sm.

Polonosporales Błaszk., Oehl, Sieverd., B.T. Goto & G.A. Silva

**Archaeosporales** C. Walker & A. Schüssler, Mycological Research 105 (12): 1418. 2001. MycoBank no.: MB 90525

Emended description: Forms symbiosis between plant roots and arbuscular mycorrhizal fungi; spore formation regularly bi-morphic, rarely only one morph or three morphs (archaeo-acaulosporoid, glomoid (sensu lato), archaeo-entrophosporoid) are known; inner walls form *de novo*; forming arbuscular mycorrhiza; vesicle formation rarely if ever described; mycelia and mycorrhizal structures do not or only faintly stain in Trypan blue, rarely darkly blue.

Type family: Archaeosporaceae J.B. Morton & D. Redecker

Other family: Antiquisporaceae G.A. Silva, Oehl, B.T. Goto, Błaszk. & Sieverd.

**Archaeosporaceae** J.B. Morton & D. Redecker, Mycologia 93 (1): 182 (2001) MycoBank no.: MB 82112

Emended description: as for the order (here above)

Type genus: *Archaeospora* J.B. Morton & D. Redecker Other genera:

*Intraspora* Oehl & Sieverd.

*Palaeospora* Oehl, Palenz., Sánchez-Castro & G.A. Silva

*Andinospora* Magurno, Uszok, Esmaeilzadeh-Salestani, Tedersoo, M.B. Queiroz & B.T. Goto

***Archaeospora*** J.B. Morton & D. Redecker, Mycologia 93 (1): 183. 2001.

MycoBank no.: MB 28457

Emended description: see Spain et al. (2003)

Type species: *Archaeospora trappei* (R.N. Ames & Linderman) J.B. Morton & D. Redecker, Mycologia 93 (1): 183. 2001.

MycoBank no.: MB 467737

Basionym: *Acaulospora trappei* R.N. Ames & Linderman, Mycotaxon 3 (3): 566. 1976.

MycoBank no.: MB 308080

***Intraspora*** Oehl & Sieverd., J. Appl. Bot. Food Qual. 80: 77. 2006.

MycoBank no.: MB 29043

Description: see Sieverding and Oehl (2006)

Type species: *Intraspora schenckii* (Sieverd. & S. Toro) Oehl & Sieverd., J. Appl. Bot. Food

Qual. 80: 77. 2006.

MycoBank no.: MB 334856

Basionym: *Entrophospora schenckii* Sieverd. & S. Toro, Mycotaxon 28: 210. 1987.

MycoBank no.: MB 129262

Synonym: *Archaeospora schenckii* (Sieverd. & S. Toro) C. Walker & A. Schüssler, The Glomeromycota: a species list with new families and new genera: 53. 2010.

MycoBank no.: MB 560049

***Palaeospora*** Oehl, Palenz., Sánchez-Castro & G.A. Silva, Nova Hedwigia 101: 92. 2015.

MycoBank no.: MB 808611

Description: see Oehl et al. (2015)

Type species: *Palaeospora spainiae* Oehl, Palenz., Sánchez-Castro & G.A. Silva, Nova Hedwigia 101: 92. 2015.

MycoBank no.: MB 547992

Synonym: *Archaeospora spainiae* (Oehl, Palenz., Sánchez-Castro & G.A. Silva) A. Schüssler & C. Walker, Mycorrhiza 29: 437. 2019.

MycoBank no.: MB 556467

***Andinospora*** Magurno, Uszok, Esmaeilzadeh-Salestani, Tedersoo, M.B. Queiroz & B.T. Goto, MycoKeys 124: 260. 2025.

MycoBank no.: MB 860983

Description: see Esmaeilzadeh-Salestani et al. (2025)

Type species: *Andinospora ecuadoriana* (A. Schüssler & C. Walker) Magurno, Uszok, Esmaeilzadeh-Salestani, Tedersoo, M.B. Queiroz & B.T. Goto, MycoKeys 124: 261. 2025. MycoBank no.: MB 860984

Basionym: *Archaeospora ecuadoriana* A. Schüssler & C. Walker, Mycorrhiza 29: 437. 2019. MycoBank no.: MB 556466

**Antiquisporaceae** G.A. Silva, Oehl, B.T. Goto, Błaszk. & Sieverd., fam. nov. MycoBank no.: MB 862462

Description: Spores acaulosporoid, formed singly in the substrate or occasionally within roots. Spores hyaline to white, small, globose to subglobose, rarely ellipsoid to ovoid. Subcellular spore structure composed of two walls: the outer wall with two hyaline layers, of which layer 1, forming the spore surface, is evanescent, and the inner wall with two permanent, flexible to semi-flexible layers. Sporiferous saccule hyaline to subhyaline, with a delicate mono- to bi-layered wall continuous with the two outer spore wall layers; usually collapsed or detached in extraradical spores. Spore walls and saccule wall staining dark in Trypan blue.

Etymology: Latin, *antiquus* (= ancient) and *spora* (= spores), referring to the phylogenetic placement of this genus within Archaeosporales, an early-diverging lineage of Glomeromycota. Type genus: *Antiquispora* Magurno, Uszok, Esmaeilzadeh-Salestani, Tedersoo, M.B. Queiroz & B.T. Goto

***Antiquispora*** Magurno, Uszok, Esmaeilzadeh-Salestani, Tedersoo, M.B. Queiroz & B.T. Goto, MycoKeys 124: 258. 2025.

MycoBank no.: MB 860980

Description: see Esmaeilzadeh-Salestani et al. (2025)

Type species: *Antiquispora disseminans* Magurno, Uszok, M.B. Queiroz & B.T. Goto, MycoKeys 124: 259. 2025.

MycoBank no.: MB 860982

**Ambisporales** Oehl, Sieverd, G.A. Silva, B.T. Goto & Błaszk., ord. nov.

MycoBank no.: MB 862460

Description: Sporocarp formation unknown; species generally bimorphic with mycorrhizal associations producing both three-walled acaulosporoid and generally mono-walled glomoid morphs on extra- or intraradical hyphae, but acaulosporoid or glomoid morph is not (yet) known for all species. *Acaulosporoid spores* produced on short hyphal appendix generally arising laterally from the hyphal neck of a terminal sporiferous saccule; with outer, middle and inner wall; germinating from the innermost wall with a germ tube emerging through the appendix attachment or penetrating the outer walls; a germination structure possibly formed between inner and middle wall. *Glomoid spores* formed singly or in loose clusters on extra- or intraradical hyphae, with a hyaline to subhyaline to creamy or brown outer layer and a hyaline to white structural layer beneath, rarely with triple walls as known for the acaulosporoid morph; germinating through the subtending hypha. Intraradical mycorrhizal hyphae, arbuscles, and vesicles stain pale blue with Trypan blue.

Etymology: Ambi- (two) -sporales: Latin ‘ambispora’, referring to the capability to produce two different spore morphs, acaulo-ambisporoid (sensu lato) and glomoid-ambisporoid.

Type family: Ambisporaceae C. Walker, Vestberg & A. Schüssler

**Ambisporaceae** C. Walker, Vestberg & A. Schüssler, Mycological Research 111 (2): 143. 2007.

MycoBank no.: MB 510208

Description: as for the order (here above)

Synonym: Appendicisporaceae C. Walker, Vestberg & A. Schüssler, Mycological Research 111 (3): 254. 2007.

MycoBank no.: MB 510505

Type genus: *Ambispora* C. Walker, Vestberg & A. Schüssler Other genera:

*Appendiculaspora* Sieverd., Oehl, Palenz. & G.A. Silva

*Ephemerapareta* G.A. Silva, Palenz., Sieverd. & Oehl *Pelotaspora* Oehl, G.A. Silva, Palenz. & Sieverd.

***Ambispora*** C. Walker, Vestberg & A. Schüssler, Mycological Research 111 (2): 147. 2007. MycoBank no.: MB 510209

Emended description: see Silva et al. (2026)

Type species: *Ambispora fennica* C. Walker, Vestberg & A. Schüssler, Mycological Research 111 (2): 148. 2007.

MycoBank no.: MB 510210

***Appendiculaspora*** Sieverd., Oehl, Palenz. & G.A. Silva

MycoBank no.: MB 862213

Description: see Silva et al. (2026)

Basionym: *Appendicispora* Spain, Oehl & Sieverding, Mycotaxon 97: 168. 2006. MycoBank no.: MB 510319

Type species: *Appendiculaspora appendicula* (Spain, Sieverd. & N.C. Schenck) Sieverd., G.A. Silva & Oehl MycoBank no.: MB 862214

Basionym: *Acaulospora appendicula* Spain, Sieverd. & N.C. Schenck, Mycologia 76: 686. 1984.

MycoBank no.: MB105884

Synonym: *Ambispora appendicula* (Spain, Sieverd. & N.C. Schenck) C. Walker, Mycological Research 112 (3): 298. 2008.

MycoBank no.: MB 511420

Synonym: *Appendicispora appendicula* (Spain, Sieverd. & N.C. Schenck) Spain, Oehl & Sieverd., Mycotaxon 97: 170. 2006.

MycoBank no.: MB 510320

***Ephemerapareta*** G.A. Silva, Palenz., Sieverd. & Oehl

MycoBank no.: MB 862218

Description: see Palenzuela et al (2011) and Silva et al. (2026)

Type species: *Ephemerapareta granatensis* (Palenz., N. Ferrol & Oehl) Oehl, Palenz., Sieverd. & G.A. Silva

MycoBank no.: MB 862219

Basionym: *Ambispora granatensis* J. Palenzuela, N. Ferrol & Oehl, Mycologia 103 (2): 334. 2011.

MycoBank no.: MB 513528

***Pelotaspora*** Oehl, G.A. Silva, Palenz. & Sieverd.

MycoBank no.: MB 862220

Description: see Silva et al. (2026)

Type species: *Pelotaspora reticulata* (Oehl & Sieverd.) Oehl, Palenz., G.A. Silva & Sieverd.

MycoBank no.: MB 862221

**Geosiphonales** Caval.-Sm., Biol. Rev. 73: 247. 1998.

MycoBank no.: MB 90546

Description: see Cavalier-Smith (1998) and Schüßler (2002)

Type family: Geosiphonaceae Engl. & E. Gilg

**Geosiphonaceae** Engl. & E. Gilg, Syllabus der Pflanzenfamilien: 24. 1924.

MycoBank no.: MB 81542

Description: see Cavalier-Smith (1998) for order, and Schüßler (2002)

Type genus: *Geosiphon* F. Wettst.

***Geosiphon*** F. Wettst., Oesterr. Bot. Z. 65 (5-6): 152. 1915.

MycoBank no.: MB 20238

Description: see Schüßler (2002)

Type species: *Geosiphon pyriformis* (Kütz.) F. Wettst., Oesterr. Bot. Z. 65 (5-6): 147. 1915.

MycoBank no.: MB 366273

Basionym: *Botrydium pyriforme* Kütz., Species algarum: 486. 1849.

MycoBank no.: MB 100748

Synonym: *Geosiphonomyces pyriformis* Cif. & Tomas., Atti Ist. Bot. Lab. Crittog. Univ. Pavia 14: 5. 1957.

**Polonosporales** Błaszk., Oehl, Sieverd., B.T. Goto & G.A. Silva, ord. nov.

MycoBank no.: MB 862461

Description: Forming hypogeous single acaulosporoid glomerospores (= spores) directly on the neck of a sporiferous saccule. Spores hyaline to white, usually globose to subglobose, with three permanent spore walls (OW, MW, IW). OW consisting of a short-lived, evanescent, thin layer, continuous with the wall of the sporiferous saccule, and a permanent, laminate, thicker layer. MW composed of one permanent, flexible to semi-flexible layer. IW permanent, coriaceous, composed of two tightly adherent layers. Only OWL2 sometimes stains reddish white in Melzer’s reagent.

Etymology: Polono- and -sporales, (English: Poland and spores), referring to the country, in which the spores of the new order were originally found.

Type family: Polonosporaceae Błaszk., Niezgoda, B.T. Goto, Magurno

**Polonosporaceae** Błaszk., Niezgoda, B.T. Goto, Magurno, Mycological Progress 20 (8): 946. 2021.

MycoBank no.: MB 840255

Description: as for the order (here above), and see Błaszkowski et al. (2021)

Type genus: *Polonospora* Błaszk., Niezgoda, B.T. Goto, Magurno

***Polonospora*** Błaszk., Niezgoda, B.T. Goto, Magurno, Mycological Progress 20 (8): 946. 2021.

MycoBank no.: MB 840256

Description: see Błaszkowski et al. (2021)

Type species: *Polonospora polonica* (Błaszk.) Błaszk., Niezgoda, B.T. Goto & Magurno, Mycological Progress 20 (8): 947. 2021.

MycoBank no.: MB 840257

Basionym: *Acaulospora polonica* Błaszk., Karstenia 27: 38. 1988. MycoBank no.: MB 133512

## Final notes

Until now all species described in the phylum Glomeromycota have been assumed to be obligate symbionts, forming arbuscular mycorrhiza with plant roots, except *Geosiphon pyriformis* that forms a symbiosis/endocytobiosis with *Nostoc*, a cyanobacteria (Gehrig et al. 1996). According to Schüßler (2002), *Ge. pyriformis* was already found in Germany and Austria, but currently this fungus is documented just in one place in the Spessart mountains of Central Germany, being probably endemic of this area. Furthermore, no environmental sequences have been found so far related to *Ge*. *pyriformis* in the NCBI. Considering the development and specific association of this fungus with cyanobacteria, we can assume that this species has a distinct ecological strategy, when compared to other glomeromycotan fungi. Probably this fact is reflected in its evolutionary history, which clearly is different from all other Archaeosporomycetes fungi, taking into consideration its unique symbiosis and endemism (Malar et al. 2021). Morphologically, glomoid spores of *Ge. pyriformis* are similar to other species in the ancestral clades of Glomeromycota (Archaeosporales and Paraglomerales, the latter belonging to Paraglomeromycetes). In the light of the specific phylogeny and ecology, we are sure that *Geosiphon* pertains to a distinct order in Archaeosporomycetes, Geosiphonales, which was described not due to its phylogeny (Cavalier-Smith 1998), but due to the huge pale asexual spores formed at hyphal tips, young hyphae without septa, and its intracellular cyanobacterial endosymbionts (Cavalier-Smith 1998). Firstly, Schüßler et al. (2001) and above all Schüßler (2002), analyzed the clear phylogenetic position of Geosiphonaceae and referred back also to the very old literature of Kützing (1849), Engler and Gilg (1924), and von Wettstein (1915).

Phylogenetically, *Geosiphon* is in a well separated clade and thus, deserving a separation of the order level. In conclusion, according to our phylogenetic trees, there is also no support to maintain the other actual families (Ambisporaceae, Archaeosporaceae, and Polonosporaceae) together in Archaeosporales. Thus, it was necessary to describe two other new orders: Ambisporales and Polonosporales. Furthermore, *Antiquispora* was described recently by Esmaeilzadeh-Salestani et al. (2025), and we have detected that this genus represents, in fact, a new family (Antiquisporaceae) in Archaeosporales, with strong support in all analyses (Figs. 1 and 2), as a sister family of Archaeosporaceae.

Another important issue to address here is related to two monospecific genera in Archaeosporaceae, *Intraspora* (Sieverding and Oehl 2006) and *Palaeospora* (Oehl et al. 2015). According to Schüßler and Walker (2010), *Intraspora* would be congeneric with *Archaeospora*. These authors affirmed, based on phylogenetic analyses, that *Intraspora schenkii* is placed among different *Archaeospora trappei* isolates and, thus, they recombined *In. schenkii* to *Ar. schenkii*. According to Bills (2015), the phylogeny indicated that *In. schenkii*, in fact, would be not congeneric, but conspecific to *Ar. trappei*. Consequently, this author proposed synonymizing this species with *Ar. trappei*. The phylogenetic analyses by Schüßler and Walker (2019) indicated that *Palaeospora spainiae* should be placed into the genus *Archaeospora* and, hence, these authors recombined *Pa. spainiae* to *Ar. spainiae*. Nonetheless, in phylogenetic analyses performed by Esmaeilzadeh-Salestani et al. (2025), it was possible to observe that sequences from different isolates of *Ar. trappei* were so dissimilar that it was possible to split them into two different clades (genera), consequently these authors described a new genus (*Antiquispora*), based on *An. disseminans* and several isolates of *Archaeospora*, beyond the already described *Archaeospora*. These authors also confirmed *Palaeospora* as a genus in Archaeosporaceae, however, they did not change the status of *Intraspora*. In our phylogenetic analyses, *Andinospora*, *Archaeospora, Intraspora*, and *Palaeospora* are placed, with strong support, in well separated clades in Archaeosporales, while *Antiquispora* represents a new family (Antiquisporaceae) within the order (Figs. 1 and 2). Furthermore, according to Silva et al. (2023; 2025), the difference in Maximum Identity (MI) for genera in the family Glomeraceae, belonging to Glomerales, is (about) 10%, considering the region “barcode” (SSU+ITS+LSU) for Glomeromycota. Last, but not least, *Intraspora* has about 90% of MI with the nearest species (*Ar. trappei*), which in fact reinforces the separation of this genus within Archaeosporaceae.

## Supporting information

Spreadsheet S1. Accession numbers for the sequences used in this study

## Acknowledgments

Daniele Magna Azevedo de Assis and Thays Gabrielle Lins de Oliveira thanks the Fundação de Amparo à Ciência e Tecnologia do Estado de Pernambuco (FACEPE) for providing fellowship. Gladstone A. Silva and Bruno Tomio Goto have a fellowship from the Conselho Nacional de Desenvolvimento Científico e Tecnológico (CNPq) (Proc. 312606/2022-2; 409181/2024-2; 306632/2022-5). Piotr Niezgoda acknowledges grant no. 2025/56/Q/NZ9/00414 from Polish National Centre of Science.

