## Supplementary material for "Revision of Archaeosporomycetes with two old and two new fungal orders: Archaeosporales, Geosiphonales, Polonosporales, and Ambisporales": Spreadsheet S1. Accession numbers for the sequences used in this study

| Species name in NCBI | ID | nrDNA region | Country | Accession numbers (nrDNA) |
|---|---|---|---|---|
| *Ambispora fennica* | Att200-23 - ex-type | SSU+ITS+LSU | Finland | FN547535, FN547536, FN547537, FN547538, FN547539, FN547540, FN547541, FN547542, FN547543, FR750157 |
| *Ambispora fennica* | Att200-21 | SSU | Finland | AM268193, AM268194, AM268195 |
| *Ambispora fennica* | Att200-23 - ex-type | SSU | Finland | AM268196 |
| *Ambispora gerdemannii* | n8_9 | SSU+ITS+LSU | Not available | JF439210 |
| *Andinospora ecuadoriana* | Att1456-7 ex-type | SSU+ITS+LSU | Ecuador | LR596344, LR596345, LR596349, LR596350, LR596351, LR596352, LR596353, LR596354 |
| *Andinospora ecuadoriana* | Att1452-6 | SSU+ITS+LSU | Ecuador | LR596347 |
| *Andinospora ecuadoriana* | Att1456-7 ex-type | SSU | Ecuador | NG070294 |
| *Andinospora ecuadoriana* | Att1452-6 | SSU | Ecuador | LR596334, LR596335, LR596336, LR596337, LR596338, LR596339, LR596340, LR596341, LR596342 |
| *Antiquispora disseminans* | Not available | SSU+ITS+LSU | Poland | PV873150, PV873151, PV873152, PV873153, PV873154, PV873155, PV873156 |
| *Antiquispora* sp. envir. sample | Not available | SSU+ITS+LSU | Not available | EUK1265821 |
| *Antiquispora* sp. envir. sample | WR668_D | SSU | New Zealand | MG829283 |
| *Appendiculaspora appendicula* | Att1235-1 | SSU+ITS+LSU | Brazil | FN547524, FN547525, FN547526, FN547527, FN547528 |
| *Appendiculaspora appendicula* | MACG1 | SSU+ITS+LSU | Peru | OM214461 |
| *Appendiculaspora callosa* | MAFF520084 | SSU+ITS+LSU | Japan | PX215129, PX215131 |
| *Appendiculaspora callosa* | MAFF520057 | SSU+ITS+LSU | Japan | PX215134, PX215135 |
| *Appendiculaspora callosa* | JA116 | SSU+ITS+LSU | Japan | PX215136, PX215137 |
| *Appendiculaspora leptoticha* | MAFF520090 | SSU+ITS+LSU | Japan | PX215138 |
| *Appendiculaspora leptoticha* | ON205A | SSU+ITS+LSU | Canada | PX215139, PX215140, PX215141 |
| *Appendiculaspora leptoticha* | MAFF520055 | SSU | Japan | AB047302 |
| *Appendiculaspora leptoticha* | NC176 | SSU | USA | AJ301861 |
| *Archaeospora europaea* | SAF115 ex-type | SSU+ITS | Switzerland | MK940274, MK940275 |
| *Archaeospora* sp. envir. sample | Not available | SSU | New Zealand | KC708349, KC708364, KC708367, KC708369 |
| *Archaeospora* sp. envir. sample | Not available | SSU | UK | JF414183 |

| | | | | |
|---|---|---|---|---|
| *Archaeospora trappei* | Att178-3 | SSU+ITS+LSU | UK | FR750034, FR750035, FR750036, FR750037, FR750038 |
| *Archaeospora trappei* | DSU-87 | SSU+ITS+LSU | Not available | ON113869 |
| *Archaeospora trappei* | NB112 | SSU | Namibia | AJ006800 |
| *Archaeospora trappei* | AU219 | SSU | Australia | AJ006801 |
| *Archaeospora trappei* | Att186-1 | SSU | Austria | Y17634, AM114274 |
| *Ephemerapareta granatensis* | ex-type | SSU | Spain | FN820272, FN820274 |
| *Geosiphon pyriformis* | GEO1 | SSU+LSU | Germany | FM876840, FM876841, FM876842, FM876843, FM876844 |
| *Geosiphon pyriformis* | GEO1 | 5.8s | Germany | AM268204 |
| *Geosiphon pyriformis* | GEO1 | SSU | Germany | Y15904, Y15905, AJ276074, AM183923 |
| *Intraspora schenkii* | Att212-4 | SSU+ITS+LSU | Argentina | FR750020, FR750021, FR750022, FR750023 |
| *Intraspora schenkii* | Not available | SSU+ITS+LSU | Not available | MZ359660 |
| *Intraspora schenkii* | CIAT -C133-8 | SSU | Colombia | FR773150 |
| *Intraspora* sp. envir. sample | WR963_H | SSU | Iceland | MG829390 |
| *Intraspora* sp. envir. sample | WR442_D | SSU | India | MG829408 |
| *Intraspora* sp. envir. sample | Not available | SSU | USA | KC708348 |
| *Palaeospora spainiae* | SAF212 ex-type | SSU+ITS+LSU | Switzerland | HG977198, HG977199, HG977200, HG977201, HG977202, HG977203 |
| *Paraglomus brasilianum* | Att260-8 | SSU+ITS+LSU | Brazil | FR750046, FR750047 |
| *Paraglomus brasilianum* | Att260-4 | SSU | Brazil | AJ301862 |
| *Paraglomus brasilianum* | BR105_2435A | SSU | Brazil | PX215163 |
| *Polonospora polonica* | ex-type | SSU+ITS+LSU | Poland | MZ359654, MZ359655, MZ359656, MZ359657, MZ359658, MZ359659 |
| *Polonospora* sp. envir. sample | Not available | SSU+ITS+LSU | Not available | EUK1265913, EUK1265914 |
| *Polonospora* sp. envir. sample | Not available | SSU+ITS+LSU | Georgia | MH982545, MH982546 |
| *Polonospora* sp. envir. sample | Not available | SSU | Not available | EUK1265901, EUK1265867 |
| *Polonospora* sp. envir. sample | G5766 | SSU | Estonia | EUK1124106 |
| *Polonospora* sp. envir. sample | G5058 | SSU | Estonia | EUK1124108 |
